# Tissue-specific genes as an underutilized resource in drug discovery

**DOI:** 10.1101/442780

**Authors:** Maria Ryaboshapkina, Mårten Hammar

## Abstract

Tissue-specific genes are believed to be good drug targets due to improved safety. Here we show that this intuitive notion is not reflected in phase 1 and 2 clinical trials, despite the historic success of tissue-specific targets and their 2.3-fold overrepresentation among targets of marketed non-oncology drugs. We compare properties of tissue-specific genes and drug targets. We show that tissue-specificity of the target may also be related to efficacy of the drug. The relationship may be indirect (enrichment in Mendelian disease genes) or direct (elevated ability to spread perturbations in human protein-protein interactome for tissue-specifically produced enzymes and secreted proteins). Reduced evolutionary conservation of tissue-specific genes may represent a bottleneck for drug projects, prompting development of novel models with smaller evolutionary gap to humans. We highlight numerous open opportunities to use tissue-specific genes in drug research and hope that the current study will facilitate discovery efforts.

---

Narrow expression in one or a few tissues is considered desirable for drug targets due to reduced risk of side effects^1,2^. Genes with narrow expression are often called ‘tissue-specific’ or ‘tissue-enriched’. Studies on microarray^3-5^ and a combination of RNA-sequencing and proteomics data^6,7^ confirm that targets of marketed drugs are biased towards tissue-specific genes. To the best of our knowledge, the first quantitative estimate was published in 2008. Dezso et al. demonstrated that tissue-specific genes are twice more likely to become drug targets than broadly expressed house-keeping genes^8^. Yang et al. confirmed a 1.7-fold higher likelihood in 2016^9^. Dezso et al. observed that tissue-specific genes may represent attractive drug targets due to their role in tissue biology and disease (e.g., brain-specific *GABRB2*, a receptor for the inhibitory neuromediator gamma-aminobutyric acid, is a target of sedative agents)^8^. These studies assessed tissue-specificity in healthy tissues. Their findings also extrapolate to diseased tissues because targets of marketed and phase 3 drugs are expressed in disease-relevant tissues even in the healthy state in 87% of the cases^10^. Also, substantial efforts are dedicated to cataloguing tissue-specific genes such as databases TiGER (2008)^11^, TiSGeD (2010)^12^, VeryGene (2011)^13^ and TissGDB (2018)^14^. Thus, systematic studies showing a significant overrepresentation of tissue-specific genes among drug targets and comprehensive resources have been available since 2008. The average time from a lead compound to entering phase 1 clinical trial is 31.2 months^15^. Let’s assume that validation of the biological hypothesis and identification of a lead compound take an equally long time. Then, ten years are sufficient for the findings of basic research to find reflection in early phase clinical trials. Now is good time to test if the industry took advantage of omics studies and pursued tissue-specific targets or not.

In this study, we examine the prevalence of tissue-specific genes among targets of marketed drugs and drugs in clinical trials. We also investigate properties of tissue-specific genes compared to drug targets. Why are these questions important to address? If tissue-specific targets are not actively pursued in early clinical trials, such study would raise awareness of open opportunities. Opportunities to discover new targets are not exhausted. A recent study by Oprea et al. indicates that only 3% of human proteins are targeted by marketed or clinical trial drugs (“Tclin”) whereas 35% have an unknown biological function and are not actively studied (“Tdark”)^16^. Also, an important parallel exists between tissue-specific genes and targets of marketed drugs. As first demonstrated in 2004, tissue-specific genes are enriched in Mendelian disorder genes^17^. The enrichment was confirmed by Yang et al. in 2016^9^. 53% targets of marketed drugs are implicated in Mendelian disorders^18^. Drugs targeting genes with a genetic link to human disease are less likely to fail in clinical trials due to lack of efficacy^19^. Thus, there may be a relationship between tissue-specificity of the target and efficacy of the drug. In fact, a recent study by Rouillard, Hurle and Agarwal concentrated on identification of omics features distinguishing targets that succeeded and failed in phase 3 trials for non-oncology diseases^20^. Phase 3 trial failures were enriched in failures due to lack of efficacy. Rouillard and colleagues limited their analysis to drugs with a single mechanism-of-action target and demonstrated that narrow expression profile of a drug target is a robust predictor of success in phase 3^20^. If we understand the relationship between tissue-specificity and efficacy and apply this knowledge to identify new, and not only tissue-specific, targets, we may reduce attrition rates in the clinic.

Here we confirm a 1.8-fold overrepresentation of tissue-specific genes among targets of marketed drugs compared to all protein-coding genes. The enrichment is 2.3-fold when non-oncology drug targets are considered separately. We observe that this historic success of tissue-specific targets is not reflected in early clinical trials neither for oncology nor for non-oncology diseases. We find two factors, that could be related to efficacy of drugs targeting tissue-specific genes. First, we confirm enrichment in disease genes among tissue-specific genes. Second, we find that tissue-specific enzymes and secreted proteins have higher ability to spread perturbations in topological analysis of human protein-protein interactome. The limiting factor for development of tissue-specific targets may be the reduced conservation of tissue-specific genes between humans and murine models and the associated challenges in preclinical research. We conclude that tissue-specific genes are a promising source for target discovery and that the translational challenges may be circumvented through creation of humanized models.

## RESULTS

Our results section is structured as follows. We investigate the prevalence of tissue-specific genes among targets of candidate and marketed drugs. Next, we explore properties that may explain depletion of tissue-specific genes among targets of drugs in early clinical trials and their overrepresentation among targets of marketed drugs. Finally, we highlight open opportunities to develop tissue-specific genes as drug targets.

We talk about genes as drug targets because the previous studies demonstrated enrichment in tissue-specific genes among drug targets based on mRNA expression^8,9^. We also define tissue-specificity based on RNA-sequencing data. We assume that the messenger RNAs are translated to their protein products, which, in turn, interact with the drugs. The concordance between gene expression and protein abundance is debated^21,22^, but a recent Ribo-seq study in rat suggests that 70 (heart) to 85% (liver) of transcribed mRNA are forwarded to translation^23^.

### Prevalence of tissue-specific drug targets

We applied peak-based definitions of tissue-specificity. We computed per-tissue Z-scores for each gene and defined tissue-specificity at nine increasingly stringent constraints: Z _second largest_ < 1/x * Z _max_, where Z_max_ denoted the Z-score in the tissue with the highest expression, Z _second largest_ denoted the Z-score in tissue with the second highest expression and x was an integer from 2 to 10 (**Fig. 1)**. Such definitions allowed genes to be expressed in multiple tissues, as long as there was a clear “peak” in the tissue with the highest expression compared to all other tissues.

**Fig. 1.**
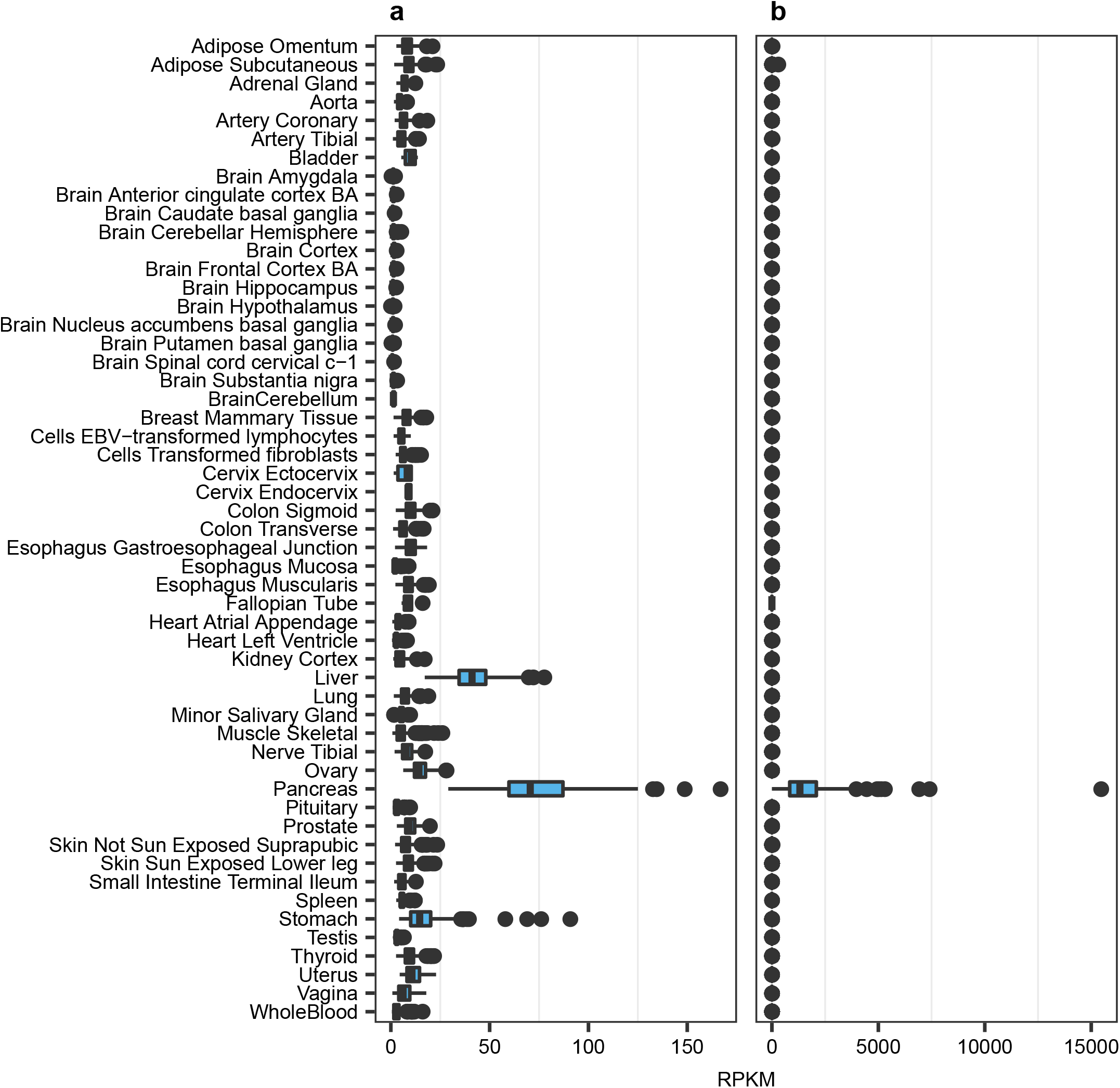
Examples of genes satisfying the most liberal (x = 2) and the most stringent (x = 10) definitions of tissue-specificity in pancreas. **a** Amino acid transporter *SLC43A1* satisfies constraint x = 2 and does not satisfy more stringent definitions. **b** Hormone insulin *INS* satisfies the most stringent definition x = 10 and consequently satisfies all more liberal definitions x = 2 to 9. The box plots show normalized expression levels in RPKM across 53 human tissues in GTEx release 6 (https://gtexportal.org/home/).

Tissue-specific genes constituted a small fraction of all human protein-coding genes (**Supplementary Data 1**). The most liberal definition x = 2 resulted in 4,573 of 18,377, 24.9% tissue-specific genes, while only 557 of 18,377, 3.0% genes satisfied the most stringent definition x = 10. If tissue-specificity was irrelevant for drug target discovery, the proportions of tissue-specific genes among drug targets would follow the ‘background’ distribution among all protein-coding genes. By contrast, we observed increasingly stronger deviations from the ‘background’ distribution with increasingly stringent definitions of tissue-specificity (**Fig. 2**). Targets of phase 1 drugs were significantly depleted of tissue-specific genes even at the liberal x = 2 (54 of 331, 16.3% < 4,573 of 18,377, 24.9%, Fisher test, p-value 1*10^−4^). Proportions of tissue-specific genes among targets of phase 2 drugs followed the ‘background’ distribution among all protein-coding genes. By contrast, targets of phase 3 drugs and marketed drugs were significantly enriched in tissue-specific genes starting from x = 6 (phase 3: 39 of 410, 9.5% > 1,018 of 18,377, 5.5%, 1.7-fold enrichment, p-value 0.001; marketed: 70 of 691, 10.1% > 1,018 of 18,377, 5.5%, 1.8-fold enrichment, p-value 2*10^−6^). Targets of withdrawn drugs were also enriched in tissue-specific genes at x = 8 to 10. The reason for withdrawal from the market was toxicity with few exceptions like unintended use for self-poisoning (barbiturates) and lack of efficacy (drotrecogin alpha). Targets of withdrawn drugs had 95% overlap (57 of 60) with targets of marketed drugs. Hence, withdrawal of these drugs from the market could not be uniquely attributed to their mechanism-of-action targets. For example, cholinergic nicotinic receptors *CHRNA1*, *CHRND* and *CHRNG* are targets of curare-like neuromuscular blocking agents. Rapacuronium bromide was withdrawn from the market due to adverse events while other drugs like vecuronium continue to be used. We used x = 6 to define tissue-specificity in all subsequent analyses, because the enrichment in tissue-specific genes among targets of marketed drugs became significant at this constraint.

**Fig. 2.**
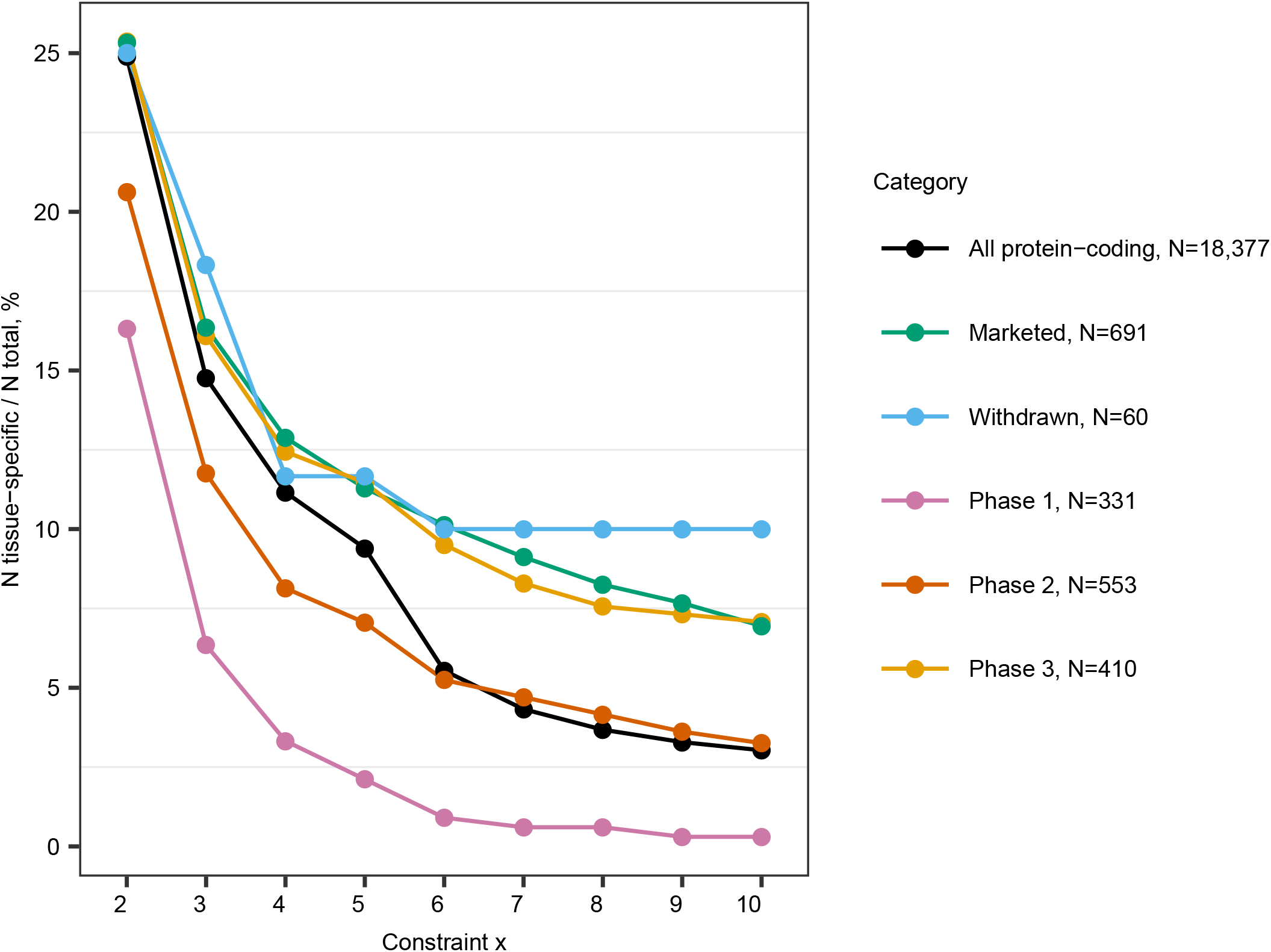
Prevalence of tissue-specific targets increased from phase 1 to the market. Percentages of tissue-specific genes among targets of drugs in each phase of clinical development were plotted in comparison to the ”background” distribution among all protein coding-genes (black line). Tissue-specificity was defined at nine increasingly stringent constraints x = 2 to 10 as illustrated in Fig. 1.

The overlap between targets of withdrawn and marketed drugs motivated us to examine ‘recycling’ of drug targets. Target genes can re-enter clinical trials when new drugs are developed for the same (e.g., generations of H_2_ histamine receptor *HRH2* blockers as anti-ulcer drugs) or a novel indication. For example, *IGF1R* is targeted by recombinant insulin growth factor 1 Mecasermin for growth failure in children (marketed agonist drug) and is evaluated as target for treatment of solid tumours (antagonist drug PL-225B in phase 1 trial NCT01779336). 431 of all 691 targets, 62.4% (**Fig. 3a**) and only 24 of 70 tissue-specific targets, 34.3% (**Fig. 3b**) were reused in clinical trials. Thus, tissue-specific genes were ‘recycled’ 1.8 times less frequently than drug targets overall (Fisher test p-value 5.6*10^−6^). Furthermore, tissue-specific genes represented an older subset of drug targets (**Fig. 3c**), although the difference was not statistically significant.

**Fig. 3.**
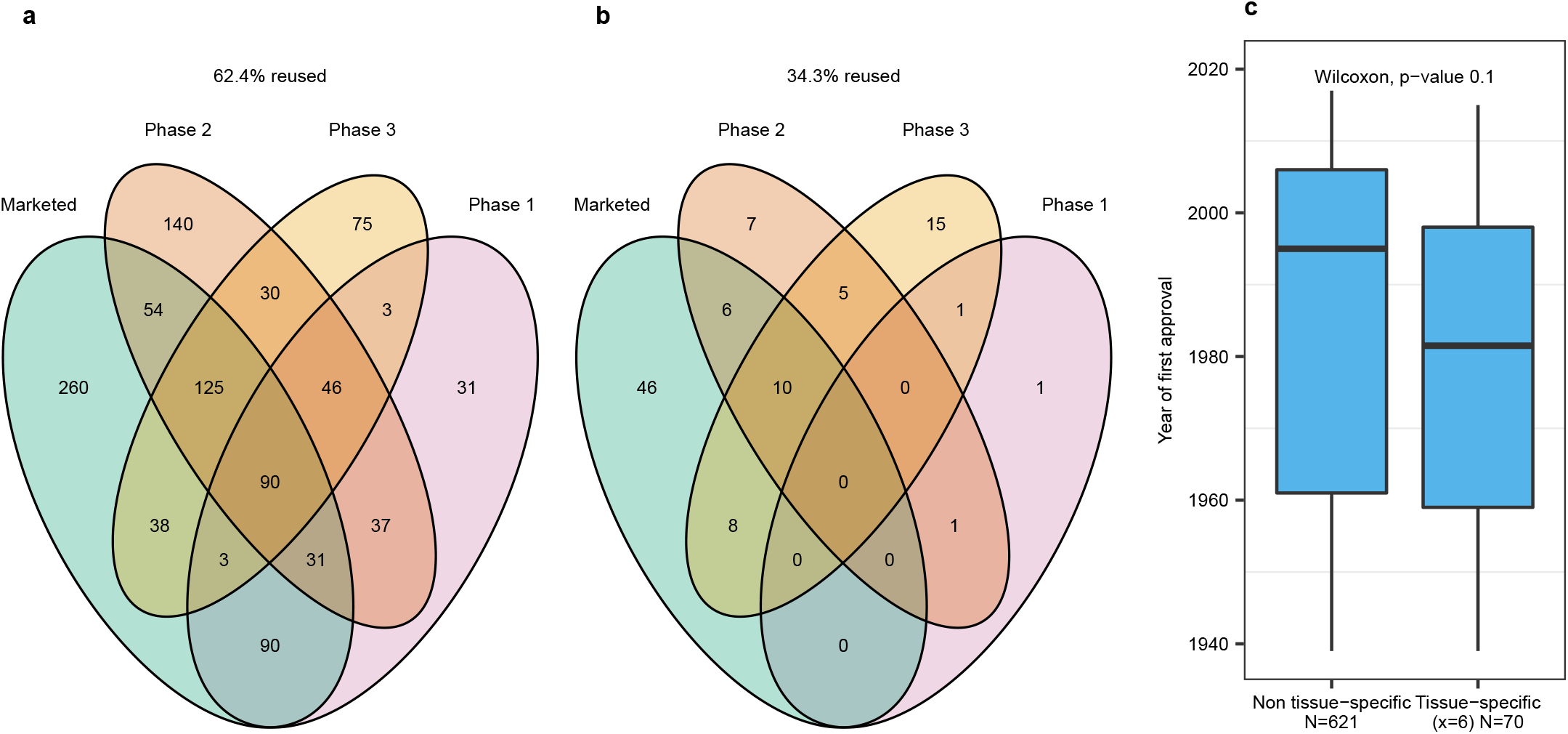
Tissue-specific targets, satisfying constraint x = 6, represented a less frequently reused (b vs a) and older (c) subset of targets of marketed drugs. **a** Reuse of all targets of marketed drugs by other drugs in clinical trials. **b** Reuse of tissue-specific (x = 6) targets of marketed drugs by other drugs in clinical trials. Venn diagrams depict the number of targets in each phase of clinical development. Overlapping areas contain genes that are targeted by several drugs in different phases of development. **c** Year of regulatory approval by FDA or another agency of the first drugs modulating non-tissue-specific targets compared to tissue-specific (x = 6) targets of marketed drugs. For example, carglumic acid was the first marketed drug modulating *CPS1* and it was approved in 2010.

In summary, tissue-specific genes, satisfying x = 6, were 1.8 times more likely to become targets of marketed drug than all protein-coding genes. However, they were not actively explored in phase 1 and 2 clinical trials. We investigated possible explanations for these trends.

### Disease indication as a confounding factor

Targets for oncology drugs are selected following different paradigms than targets for non-oncology drugs. For example, traditional cytotoxic agents aim to induce cell death or inhibit growths through core cell processes, that are carried out by ubiquitously expressed targets like DNA topoisomerase II (etoposide). Oncology drugs have different safety profiles from non-oncology drugs, with more side effects being tolerated. Also, some drugs target cancer-specific mutant proteins, which are not captured by gene expression analysis on healthy tissue. For example, vemurafenib targets mutated *BRAF* in melanoma according to the FDA label (www.accessdata.fda.gov/drugsatfda_docs/label/2017/202429s012lbl.pdf).

Targets of phase 1 drugs were predominantly investigated for oncology indications: 253 of 331, 76,4%. Targets of phase 2 drugs displayed an almost balanced representation of targets for oncology - 239 of 553, 43.2% - and non-oncology indications - 314 of 553, 56.8%. By contrast, most targets of phase 3 drugs - 265 of 410, 64.6% - and of marketed drugs – 479 of 691, 69.3% - were developed for non-oncology indications. Prevalence of tissue-specific genes among targets of clinical trial and marketed drugs was confounded by disease indications.

Hence, we examined oncology and non-oncology targets separately (**Supplementary Fig. 1**). Targets of phase 1 drugs were depleted of tissue-specific genes irrespective of disease indication. The discrepancies between oncology and non-oncology targets started to emerge in phase 2. Targets of marketed non-oncology drugs displayed a 2.3-fold overrepresentation in tissue-specific genes (at x = 6: 61 of 479, 12.7% > 1,018 of 18,377, 5.5%, Fisher test, p-value 3.6*10^−9^), which was stronger compared to pooled analysis for all disease indications. By contrast, tissue-specific genes were underrepresented among targets of oncology drugs.

### Insights from evolutionary biology and population genetics

Evolutionary properties may explain the underrepresentation of tissue-specific targets in early clinical trials. Wenhua Lv et al. demonstrated that targets of FDA-approved drugs are more evolutionary conserved than non-target genes^24^. By contrast, **i**n 2004, Winter, Goodstasdt and Ponting investigated expression of 4,960 human genes in 27 tissues and demonstrated that tissue-specific genes are less evolutionary conserved than broadly expressed genes using K_a_/K_s_ ratios^17^. To clarify, K_a_/K_s_ is the rate of nonsynonymous K_a_ to synonymous K_s_ amino acid changes in a pair of orthologs. Low K_a_/K_s_ implies that nonsynonymous changes are selected out, while K_a_/K_s_ exceeding 1 may indicate that changes are favored and retained as in immune genes adapting to new pathogens^25^.

We revisited the analysis with the current larger data set. We examined K_a_/K_s_ ratios for human protein-coding genes and their mouse counterparts because mice are the most common species in preclinical research. We confirmed opposite patterns for K_a_/K_s_ ratios of tissue-specific genes and drug targets (**Fig. 4a**). Tissue-specific genes had significantly higher K_a_/K_s_ than all protein-coding genes (Mann-Whitney U test, Bonferroni adjusted p-value 1*10^−40^). By contrast, targets of marketed and clinical trial drugs were significantly more conserved (the highest Bonferroni adjusted p-value was 1*10^−5^ for oncology drug targets in phase 3). The trend held for targets for oncology and non-oncology indications. K_a_/K_s_ ratios were inversely correlated with sequence identity (Spearman rho -0.86) and similarity (-0.82) between human proteins and their mouse orthologs. Conservation of protein sequence is considered a proxy for conservation of biological function^26^. Also, 283 of 1,018 tissue-specific genes (27.8%) compared to 2,719 of all 18,377 protein-coding genes (14.8%) did not have a unique ortholog in mouse. Therefore, absence of a convenient animal model and gaps in translation from animal research to clinical trials in humans may complicate development of tissue-specific genes as drug targets.

**Fig. 4.**
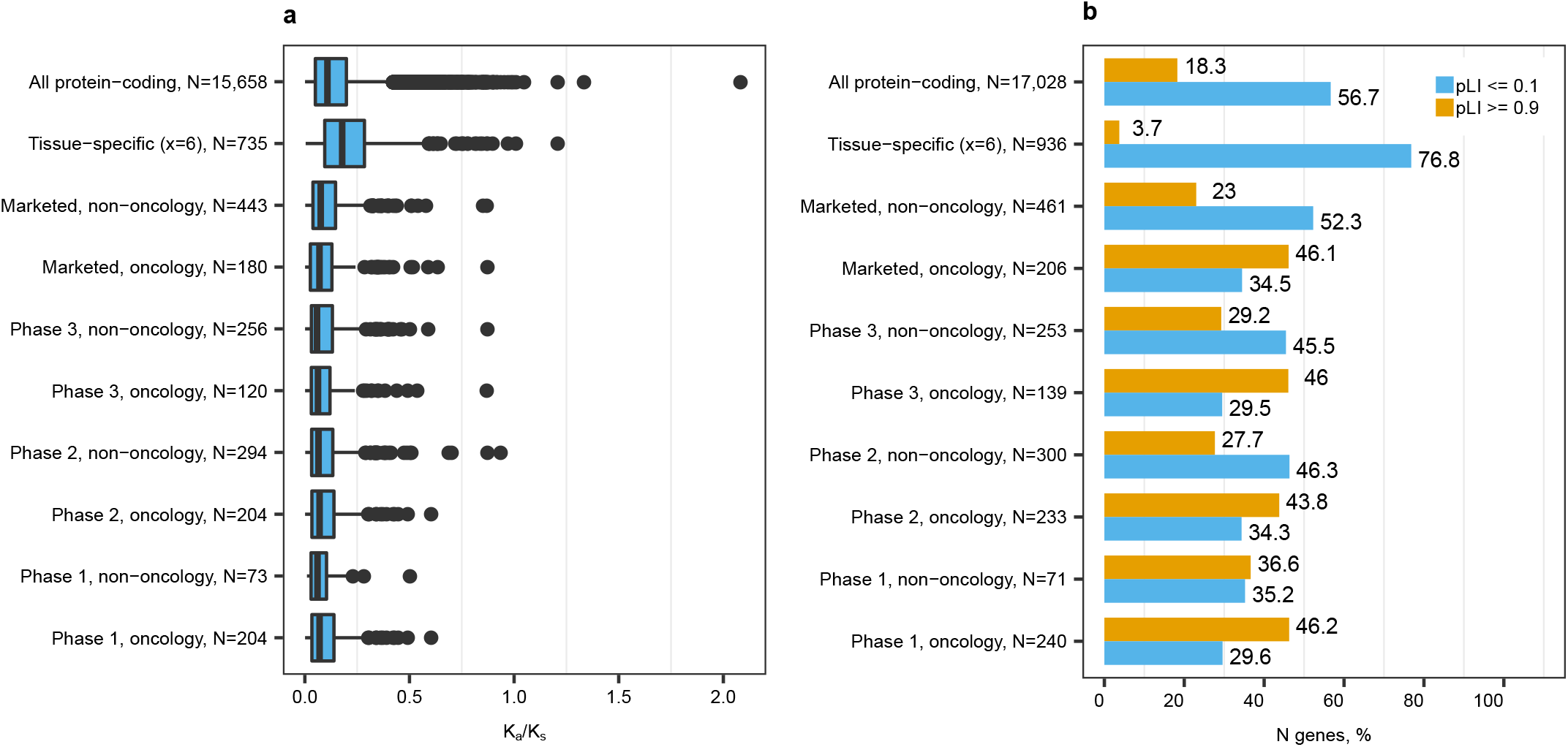
Tissue-specific genes were subject to less strong selection pressure compared to all protein-coding genes and drug targets. **a** Ka/Ks ratios for human-mouse 1:1 orthologs. 1:1 ortholog refers to a human gene with one unique counterpart in mouse as opposed to 1-to-many or many-to-many orthologs that arise from duplication or gene fusion events. **b** Percentages of loss-of-function intolerant (ExAC consortium pLI *≥* 0.9) and tolerant (pLI *≤* 0.1) genes among tissue-specific genes and drug targets compared to all protein-coding genes. Discrepancies in sample size are due to different numbers of genes mapped to the respective data sets.

We next examined selection pressure within the human species using a new metric, that was recently developed by the ExAC consortium - probability of being loss-of-function intolerant (pLI)^27^. Genes with high pLI have significantly lower observed than expected frequencies of loss-of-function variants, indicating that deleterious variants in these genes are selected out of the human population. Genes with pLI >= 0.9 are considered loss-of-function intolerant and their “knockout” in humans implies “some non-trivial survival or reproductive disadvantage”^27^. By contrast, genes with pLI <= 0.1 are considered loss-of-function tolerant^27^. In our analysis, tissue-specific genes were enriched in loss-of-function tolerant and depleted of intolerant genes compared to all protein-coding genes (**Fig. 4b**). By contrast, targets of oncology drugs were enriched in loss-of-function intolerant (highest Bonferroni adjusted p-value 8*10^−13^ in phase 3) and depleted of tolerant genes (highest Bonferroni adjusted p-value 3*10^−9^ for marketed drugs). Targets of marketed non-oncology drugs had comparable prevalence of loss-of-function tolerant and intolerant genes compared to all protein-coding genes. The distributions of pLI confirmed that tissue-specific genes were more likely to become targets for non-oncology drugs.

Genes with pLI >= 0.9 are more likely to be detected in genome-wide association studies (GWAS)^27^, and to attract attention as candidate drug targets through GWAS. We investigated whether less conserved tissue-specific genes were less frequently found in GWAS. GWAS variants are often located in intergenic regions and can be mapped to candidate genes by proximity on the chromosome or through an association between genotype of the GWAS variant and expression of a gene (eQTL). Mapping through eQTL can highlight regulatory relationships in disease-relevant tissues^28^, and, consequently, is frequently used. The two types of mapping can highlight different candidate genes^29,30^, and require follow-up experiments to determine causal genes. Tissue-specific genes were equally likely to be detected as a nearest gene to a GWAS variant but 1.3 times less likely to be mapped from GWAS to single-tissue cis-eQTLs than all protein-coding genes (Fisher test, Bonferroni adjusted p-value 2*10^−9^). Interestingly, only mapping by proximity on chromosome distinguished drug targets from all protein-coding genes (**Supplementary Fig. 2**). 111 of 392 (28.3%) of GWAS to eQTL relationships for tissue-specific genes were detected in the corresponding tissues with highest expression. These results were not surprising. Our definition of tissue-specificity allowed lower expression in other tissues. Some tissues, including kidney cortex with 48 tissue-specific genes, had no eQTL data due to insufficiently high number of samples and did not contribute to this analysis. Also, approximately a third of GWAS to eQTL relationships can only be captured using multiple tissues, while single-tissue analyses lack power to detect the associations^30^. Thus, tissue-specific genes were less likely to be highlighted as candidate targets if investigators relied on the GWAS to eQTL approach.

In summary, underrepresentation of tissue-specific targets in early clinical trials could be attributed to their primary relevance for non-oncology diseases and translational challenges. Despite these challenges, tissue-specific genes were enriched among targets of marketed drugs. We hypothesized that tissue-specificity was related to efficacy and not only to safety.

### Tissue-specificity vs efficacy

Efficacy of a drug can be viewed as a combination of properties of the drug (e.g., potency, bioavailability, selectivity etc.) and properties of its intended target(s). Here, we focus on efficacy-related properties of the targets.

#### Prevalence of disease genes

Drugs, that modulate targets with genetic evidence for a human disease, are less likely to fail in clinical trials for lack of efficacy^19^. Knowledge of human genetics can help to understand the biological function of the target, find target engagement biomarkers for clinical trials and estimate dose-response curves^31^. These factors can enhance the chances of a drug to succeed.

We compared the prevalence of OMIM^32^ and Protein Truncating Variants *esc*aping nonsense mediated decay (PTVesc) genes^33^ among tissue-specific genes and drug targets. OMIM genes have an entry in the Online Mendelian Inheritance in Man^®^ Morbid Map data base^32^, are well-known disease genes and are likely to be explored in target discovery. By constrast, PTVesc genes are an emerging class of candidate genes that can cause disease by gain-of-function mechanism. PTVesc genes are significantly depleted of genetic variants, that result in mRNA that escape nonsense-mediated decay and production of truncated proteins with altered function (e.g., *PNPLA3* and *APOL1*)^33^. Methods for detection of PTVesc are recently developed, so PTVesc genes are unlikely to be explored to the same extend as OMIM genes. Tissue-specific genes were enriched in both OMIM and PTVesc genes (**Fig. 5**). The outcomes of Fisher exact test for tissue-specific genes were 272 of 1,018 > 3,870 of 18,377, Bonferroni adjusted p-value 2*10^−4^ for OMIM genes and 159 of 1,018 > 1,913 of 18,377, Bonferroni adjusted p-value 4*10^−6^ for PTVesc genes. As expected, drug targets for oncology and non-oncology indications across all phases of clinical development were enriched only in OMIM genes (**Fig. 5a**). By contrast, the prevalence of PTVesc genes among drug targets did not significantly deviate from the overall prevalence among protein-coding genes (**Fig. 5b**).

**Fig. 5.**
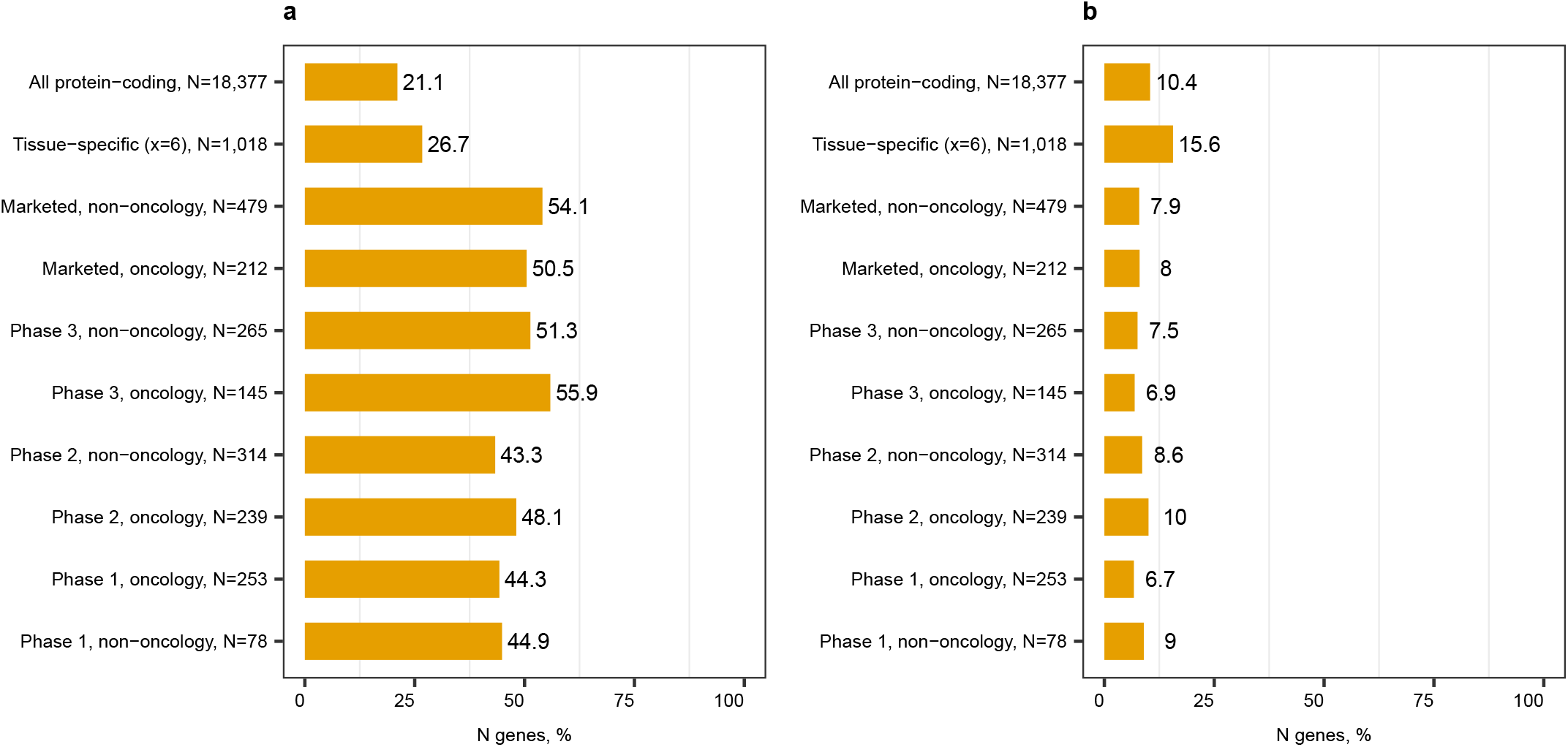
Tissue-specific genes were enriched in disease genes and potential disease genes with gain-of-function mechanism. The bars show percentages of **a** OMIM and **b** PTVesc genes in each gene category.

In total, 386 of 1,018 tissue-specific genes (37.9%) were OMIM genes or PTVesc genes or both. Thus, tissue-specific genes were more likely to provide necessary information for development of efficacious drugs through human genetics than protein-coding genes overall.

#### Network analysis

The ability to spread perturbations through the cell and cause phenotypic changes is a key property of drug targets, which is reflected by topological properties in protein-protein interaction (PPI) networks^34^. We explain the network topology properties in **Supplementary Fig. 3**. We recommend section 2 in^35^ for a detailed explanation of the relationship between network topology properties and spread of perturbations. We performed network analysis on STRING v10.5^36^ (**Supplementary Data 2**) because tissue-specific proteins are well represented in this data base^37^. We included three gene sets with known ability to affect phenotype as controls. Distribution of network-topological properties of these genes should indicate if the PPI network accurately reflects the ability of genes to spread perturbations and cause phenotype. Essential genes cause cell death or hamper growth upon silencing in two human cancer cell lines^38^. These genes serve as positive control for severe phenotypes. OMIM genes cause disease and serve as positive control for less severe phenotypes. Genes with rare homozygous loss-of-function rhLOF variants in three human cohorts serve as negative no-phenotype controls (British-Pakistani, ExAC and Icelandic individuals, Suppl. Table 2 from^39^). The human subjects come from the general population and are assumed to be healthy, so loss of function of rhLOF genes is assumed to be compensated. No association between presence of rhLOF genes and rate of drug prescriptions and medical consultations has been confirmed in the British-Pakistani cohort^39^.

First, we investigated the sources of supporting evidence for PPIs (**Supplementary Fig. 4**). Tissue-specific genes did not markedly differ from all protein-coding genes in this respect. Each PPI had a score reflecting the amount of cumulative evidence supporting existence of the interaction. Interestingly, non-oncology drug targets from phase 1 to the market tended to have more high confidence PPIs than other gene categories (**a**). PPIs for oncology drug targets and essential genes had more support from co-expression across multiple experiments and tissues (**f**) and the experimental evidence channel (**g**). PPIs of non-oncology drug targets tended to have more support from pathway data bases (**h**). We concluded that indirect (*functional*) interactions were important for non-oncology targets and kept both physical and functional interactions for analysis. Most PPIs were supported by published scientific literature (**i**). The number of reported PPIs and the number of published articles per gene were correlated (Kendall tau b = 0.31), indicating a source of bias for network topology analysis. The neighbourhood (**c**), fusion (**d**) and co-occurrence (**e**) channels provided support for relatively few PPIs, consistent with primary relevance of these three evidence channels for PPIs in Archaea and Bacteria^36^.

We observed that most PPIs had low confidence scores even for drug targets (**b**). Mora and Donaldson demonstrated that removing low confidence interaction does not substantially improve the ability to discriminate drug targets based on their topological properties^40^. Hence, we analysed the complete interaction set, but directly incorporated the confidence in PPIs into the calculations and computed weighted topological properties (see **Methods/Network analysis** for details). The calculations were performed on the largest connected component including 19,574 proteins and 5,676,527 PPIs. The network diameter (unweighted) was 6. Topological properties of the nodes accurately reflected their ability to spread perturbations through the network (**Fig. 6** and **Table 1**). Distributions of centrality scores among rhLOF genes did not significantly differ from the overall distributions among protein-coding genes (except for slightly lower closeness centrality scores). Drug targets, OMIM genes and essential genes had elevated centrality scores. Betweenness centrality was the only topological property that could distinguish tissue-specific genes from all protein-coding genes (**Table 1**). The trend was nominally significant but did not pass the correction for multiple testing. Our results were consistent with the previous study on regulatory networks, in which the Sonawane et al. applied a less stringent definition of tissue-specificity and found that tissue-specific genes serve as “bottlenecks” on signaling paths^41^.

**Fig. 6.**
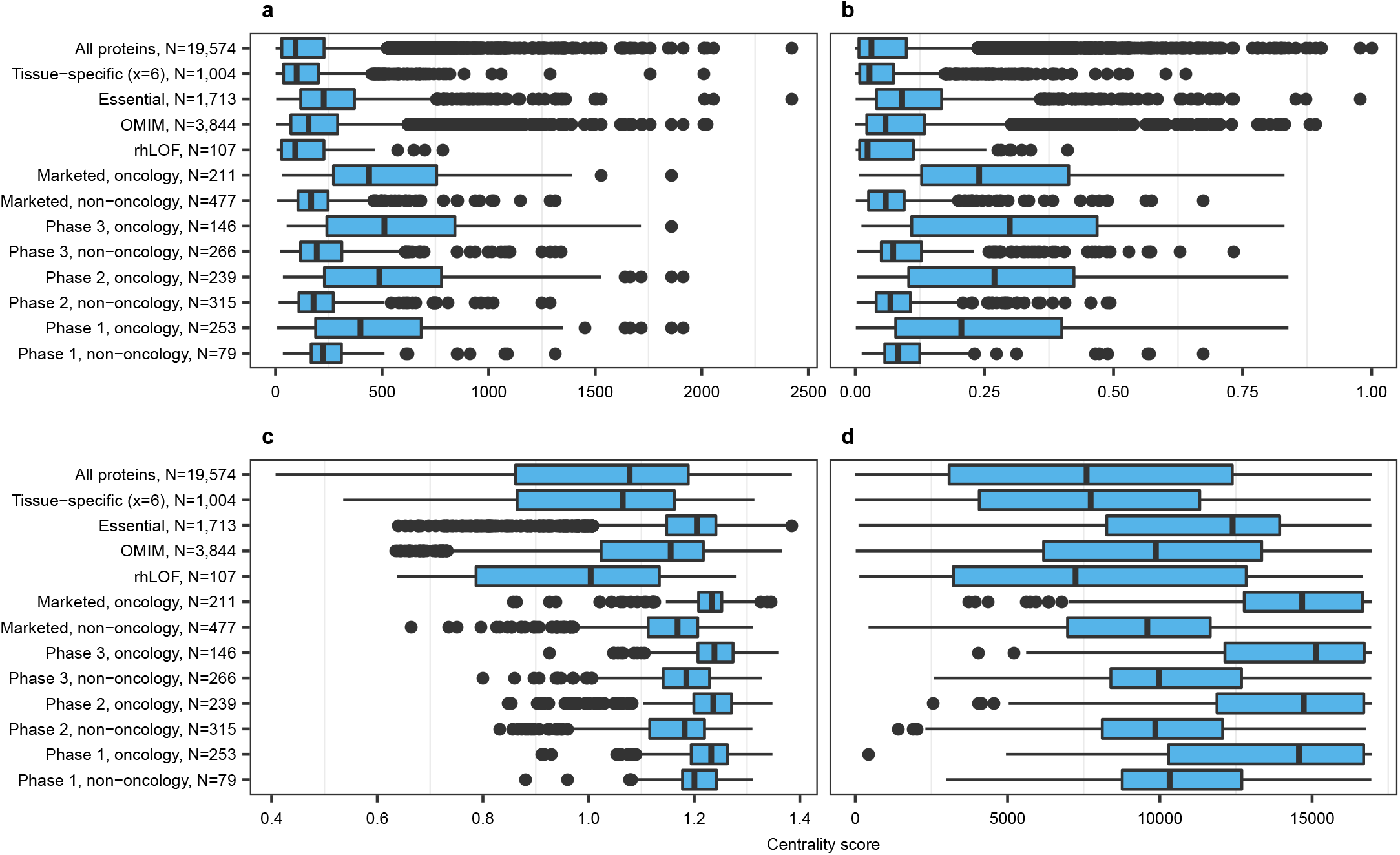
Centrality scores in STRING v 10.5. **a** Strength **b** Eigenvector centrality **c** Closeness centrality (normalized) **d** Weighted k-shell. Discrepancies in sample size are due to different numbers of genes mapped between data sets.

**Table 1.** Betweenness centrality scores in STRING v 10.5. Betweenness centrality scores are displayed separately in tabular form due to skewed distributions. IQR stands for interquartile range. P-values are from two tailed Mann-Whitney U test between the gene categories and all protein-coding genes (marked as ‘Reference’).

| Group | Median (IQR) | Nominal p | Bonferroni p |
| --- | --- | --- | --- |
| All proteins, N=19,574 | 14 (0-19,548) | Reference | Reference |
| Tissue-specific (x=6), N=1,004 | 32.1 (0-19,536.3) | $7.3 \times 10^{-3}$ | 0.088 |
| Essential, N=1,713 | 29,414.2 (40-130,570) | $8.9 \times 10^{-179}$ | $1.1 \times 10^{-177}$ |
| OMIM, N=3,844 | 14,754.6 (7-59,036.2) | $2.6 \times 10^{-169}$ | $3.1 \times 10^{-168}$ |
| rhLOF, N=107 | 8 (0-2,241.5) | 0.2 | 1 |
| Marketed, oncology, N=211 | 48,734.4 (11,077.1-201,844.9) | $2.0 \times 10^{-46}$ | $2.4 \times 10^{-45}$ |
| Marketed, non-oncology, N=477 | 1,331.4 (13-44,616.7) | $5.5 \times 10^{-23}$ | $6.6 \times 10^{-22}$ |
| Phase 3, oncology, N=146 | 69,496.3 (12,503.2-357,208.5) | $3.6 \times 10^{-33}$ | $4.3 \times 10^{-32}$ |
| Phase 3, non-oncology, N=266 | 8,921.6 (38.5-59,684.4) | $2.7 \times 10^{-21}$ | $3.3 \times 10^{-20}$ |
| Phase 2, oncology, N=239 | 58,807 (595.9-375,702) | $9.4 \times 10^{-46}$ | $1.1 \times 10^{-44}$ |
| Phase 2, non-oncology, N=315 | 1,996.8 (9-53,038) | $2.5 \times 10^{-15}$ | $3.0 \times 10^{-14}$ |
| Phase 1, oncology, N=253 | 51,848.7 (5,140.7-222,555.3) | $6.8 \times 10^{-46}$ | $8.1 \times 10^{-45}$ |
| Phase 1, non-oncology, N=79 | 19,390.9 (53.2-7,7431.6) | $1.4 \times 10^{-7}$ | $1.7 \times 10^{-6}$ |

We further investigated which tissue-specific genes had high betweenness centrality scores. The ten highest betweenness centrality scores were for genes encoding hormones (insulin *INS*; glucagon *GCG*; *POMC* giving rise to adrenocorticotrophin and lipotropin beta in the pituitary), other secreted proteins (albumin *ALB*; neuropeptide S *NPS*; plasminogen *PLG*; *APOA1*, a major constituent of high density lipoprotein cholesterol), rate limiting enzyme in synthesis of bile acids *CYP7A1*, mitochondrial enzyme *FDXR* and electron transporter *FDX1* acting together in synthesis of steroid hormones in the adrenal glands. Enzymes and secreted proteins, that were expressed in a tissue-specific manner, had higher betweenness centrality scores than other tissue-specific genes (**Supplementary Table 1**). These genes may have important *functional* interactions and their modulation may cause effects outside of their tissue-of-origin. For example, aliskiren fumarate inhibits the kidney-specific enzyme renin *REN* that is part of the renin–angiotensin–aldosterone system, lowers blood pressure and mitigates manifestations of hypertension in the whole body.

### Historic precedents to guide future applications

In total, only 100 of 1,018 (9.8%) tissue-specific genes were explored as targets of marketed or clinical trial drugs. 284 of the remaining 918 (30.9%) tissue-specific genes were classified as Tdark in the TCRD data base^42^, i.e., were poorly researched with unknown biological function. 529 of 918 (57.6%) showed some indication of druggability by small molecule or antibody approaches (**Supplementary Fig. 5**). The definition of druggability constantly expands, and targets that cannot be modulated with small molecules or antibodies may be targeted by antisense oligonucleotides or other approaches. Hence, the opportunities to identify novel drug targets among tissue-specific genes were not exhausted. In the next subsections, we review how tissue-specific genes have been used historically and highlight promising future applications.

#### Tissue-specific drug targets

Tissue-specific genes are targeted by drugs approved for diverse disease indications (**Table 2**). Tissue-specific genes can be targeted by small molecules (e.g., *ACE* - captopril), analogues of endogenous substances (*AVPR1B* – desmopressin acetate, an analogue of vasopressin), antibodies (*TNF* - etanercept) and new modalities. We and other researchers^9,17^ demonstrated that tissue-specific genes are enriched in OMIM genes. Hence, defective forms of tissue-specific genes causing rare monogenic diseases are potential targets for genome editing with the emerging CRISPR/Cas9 technology (e.g., surfactant genes in surfactant deficiencies^43^, *SERPINA1* in alpha-1 antitrypsin deficiency^44^).

**Table 2.**
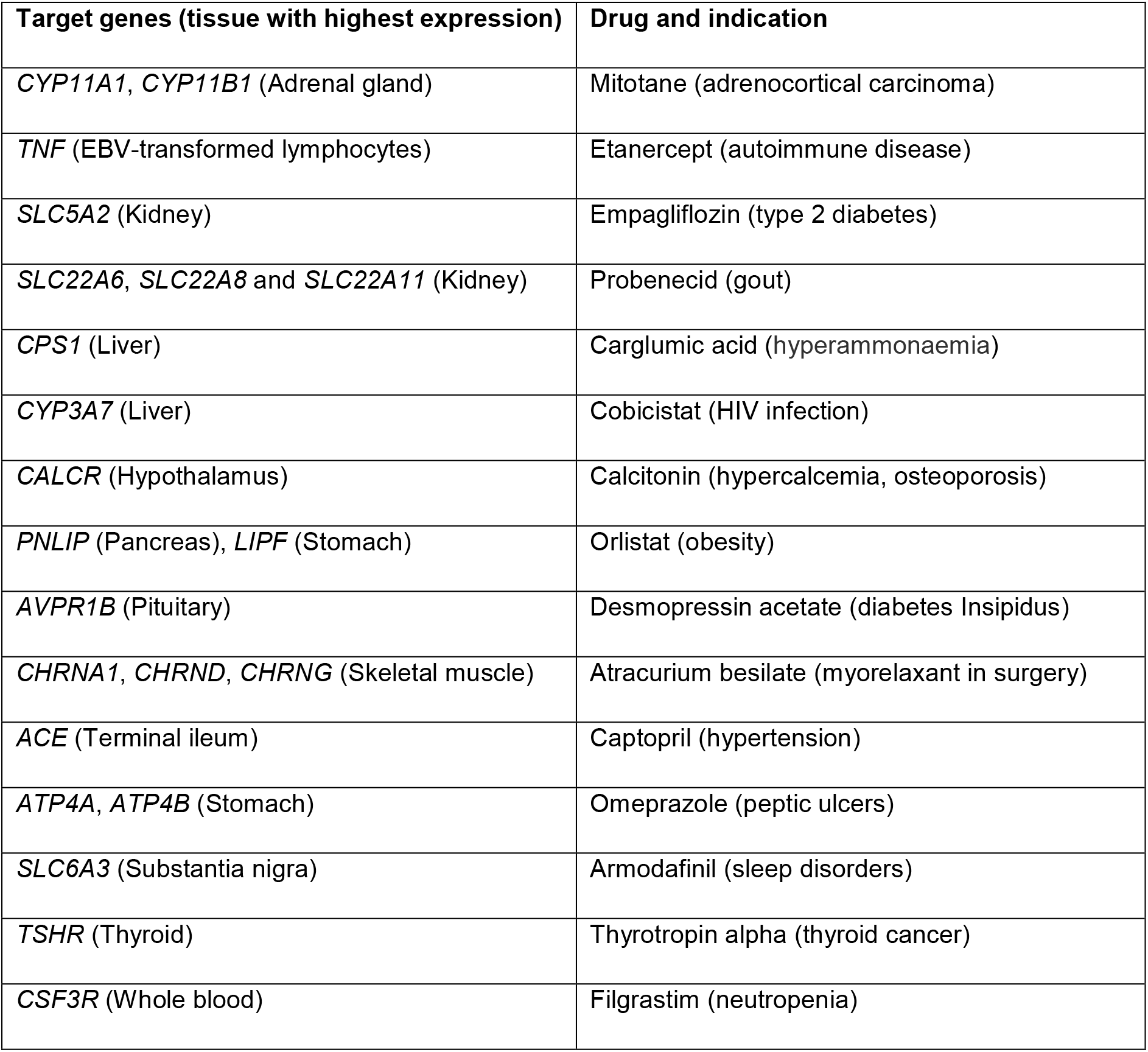
Examples of tissue-specific (x = 6) targets of marketed drugs.

Historically, tissue-specific genes were predominantly targets for non-oncology diseases. However, tissue-specific genes also find applications in oncology (e.g., mitotane for endocrine therapy in inoperable adrenocortical carcinoma^45^) and as targets for pharmacoenhancers. These application scenarios may be expanded in the future.

The pharmacoenhancer Cobicistat is administered together with antiretroviral drugs and inhibits cytochromes of *CYP3A* subfamily that degrade antiretroviral drugs primarily in the liver and intestines. Cobicistat helps to maintain therapeutic concentration of antiretroviral agents for a longer time whereby improving adherence to therapy in HIV patients^46^. Drug-metabolizing enzymes such as the liver-specific cytochromes *CYP1A2*, *CYP2D6* and *CYP2C9^47^* may be candidate targets for other pharmcoenhancers to improve bioavailability or prolong action of the main drug.

Tissue-specific genes represent potential targets for antibody-based therapies (e.g., mammary gland specific transcription factor *ANKRD30A* for breast cancer^48^). Promoters of tissue-specific genes can be used in oncolytic viral therapies to achieve specific expression of the virus in the target tissue (e.g., urothelium-specific adenovirus CG8840 with uroplakin 2 *UPK2* promoter for bladder cancer^49^ and prostate-specific antigen *KLK3* targeted adenovirus CG7870 for prostate cancer^50^). N-acetylgalactosamine (GalNAc)-conjugated antisense oligonucleotide drugs bind to the liver-specific *ASGR1* and enable targeted delivery to hepatocytes^51^. Similarly, tissue-specific genes can be explored as targets for other targeted delivery approaches, and not only in cancer.

Finally, *TNF* exemplifies a category of genes that dramatically change their expression in disease and become specific to inflamed or cancerous tissue (represented by EBV-transformed lymphoblastoid cell line and transformed fibroblasts in GTEx). Anti-TNF drugs like etanercept are used to treat rheumatoid arthritis, psoriatic arthritis and ankylosing spondylitis. *TNF* is 3.55 log_2_ fold (11.7 times on linear scale) higher in synovial membranes of recently diagnosed patients with psoriathic arthritis, who are naïve to anti-TNF treatment, than in healthy donors^52^. This example highlights the importance of extensive tissue panels including both healthy and disease tissues such as E-MTAB-3732^53^ for target discovery in cancer and inflammatory diseases.

#### Other applications

Tissue-specifically produced secreted proteins (or the corresponding recombinant peptides, proteins and other synthetic analogs) are used as replacement therapy. The best-known examples include hormone replacement therapies (insulin in type 1 diabetes, thyroid hormone in hypofunction of thyroid gland, oxytocin to induce labour, etc.) and medication containing pancreatic or gastric enzymes to aid digestion (e.g., Creon). Other replacement therapies are in development. For example, FDA recently approved the coagulation factor-albumin fusion protein Idelvion for congenital complement factor IX *F9* deficiency while other complement replacement therapies are in clinical trials^54^. In our analysis, 268 of 918 tissue-specific genes, that were not yet explored as targets of marketed or clinical trial drugs, encoded secreted proteins, which may represent opportunities for new replacement therapies (e.g., hormones and their combinations for treatment of type 2 diabetes and obesity^55^, artificial saliva with recombinant lysozyme in treatment of xerostomia and Sjögren’s syndrome^56^).

Tissue-specific genes with protein products entering the bloodstream can serve as biomarkers to monitor the state or function of a tissue. For example, *KLK3*, better known as prostate cancer antigen, has prostate-specific expression, enters bloodstream, displays elevated levels in prostate cancer and benign hyperplasia of the prostate and is used as a pre-screening test for prostate cancer^57^. Tissue-specific biomarkers indicating tissue damage have several conceptual advantages over conventional laboratory tests including higher specificity^58^. For example, circulating proteins with liver-specific expression are evaluated as markers of acetaminophen induced hepatotoxicity^59^. Biomarkers discussed in literature tended to conform to more liberal definitions of tissue-specificity, which may be sufficiently stringent for successful development of biomarkers (e.g., *RBP4*, evaluated in^59^, satisfied constraint x = 5 in liver and had lower expression in adipose tissues and pituitary, *NPPB* encoding NT-pro-BNP satisfied x = 4 in heart atrial appendage with high expression in some left ventricle samples).

#### Non-coding genes

We focused on protein-coding genes, but long non-coding RNA (lncRNA) also tend to be expressed in tissue- and cell-type specific manner^60^. We identified 2,113 long non-coding RNAs (**Supplementary Data 3**), from which 77 were tissue-specific at x = 6 and confirmed polyadenylated, i.e., reliably detected with the GTEx RNA sequencing protocol. Long non-coding RNAs have potential as drug targets^61^. For example, prostate-specific lncRNA *PCGEM1* is candidate target in prostate cancer^62^.

## DISCUSSION

We conducted a retrospective analysis of tissue-specific genes compared to drug targets in all phases of clinical development. Targets of phase 1 drugs reflect the most recent research. By contrast, targets of marketed drugs have undergone at least a decade in preclinical and clinical development and reflect older research. Theoretically, overrepresentation of tissue-specific targets on the market and depletion in phase 1 could reflect a historic shift in target selection paradigms. However, Rouillard and colleagues^20^ studied phase 3 drugs (projects with comparable “age”) and demonstrated that drugs modulating tissue-specific targets are more likely to succeed in phase 3 and gain regulatory approval. Thus, the data presented in Fig. 2 and Supplementary Fig. 1 do not merely represent a historic trend. We are justified to state that drugs modulating tissue-specific targets are indeed more likely to progress in the clinic. We observed that tissue-specific genes, satisfying x = 6, were 1.8 times more likely to become targets of marketed drugs and 2.3 times more likely to become targets of marketed non-oncology drugs than protein-coding genes overall. Our findings were consistent with the previous studies^8,9^.

Success of tissue-specific genes as drug targets may be due to a complex combination of factors. Good understanding of target biology is essential for target-based drug discovery. Tissue-specific genes are involved in specialized tissue functions and human diseases, in which genetic evidence can provide the necessary supporting information for development of new drugs. By contrast, broadly expressed genes may have diverging functions and protein isoforms with distinct subcellular localization in different tissues^7^. The biology of tissue-specific genes may be less complex to study than the biology of broadly expressed genes, which may contribute to the development of efficacious drugs. The ability to spread perturbations within and outside the tissue-of origin may be a direct contributing factor to efficacy of drugs modulating tissue-specific genes with high betweenness centrality scores, especially enzymes and genes encoding secreted proteins. Narrow expression profile decreases the probability of side effects in non-intended target tissues. Tissue-specific genes are depleted in loss-of-function intolerant genes, have average number of neighbors and are located in interactome regions with average connectivity as indicated by distributions of strength and eigenvector centrality scores. These properties indicate improved safety, as they are in sharp contrast to oncology targets, whose modulation can cause severe side effect.

Feasibility is another crucial component to the success of a drug program. Murine models are commonly used to assess toxicity and for early efficacy studies *in vivo* and can serve as a ‘filter’ to make stop/go decisions for a drug project. The targets of drugs from phase 1 to the market are biased towards evolutionary conserved genes. We suggest using humanized mouse models or other non-rodent species with smaller evolutionary distance to humans to overcome translational challenges and enable the development of tissue-specific genes and other less conserved biological entities like long non-coding RNAs^63^ as drug targets.

## MATERIALS AND METHODS

### Gene expression

Gene-level RPKM values were downloaded from The Genotype-Tissue Expression Consortium^30^ (https://gtexportal.org/home/, release 6). The per-tissue mean RPKM for each gene was subjected to Z-transformation across tissues and then to a second Z-transformation across genes to bring all Z-scores to the same scale. We identified 18,377 protein-coding genes and 2,113 long non-coding RNA with HGNC approved gene symbol^64^. The non-alternative loci data set was obtained from the HGNC Database (www.genenames.org, 30.08.2017).

### Drug targets

Mechanism-of-action targets of marketed and clinical trial drugs, disease indications and year of first approval for marketed drugs were extracted from ChEMBL version 23^65^. Drugs were classified as phase 1, 2, 3 or marketed drugs based on the maximal phase they reached in clinical trials. Disease indications were mapped to Disease Ontology^66^. Proteins were classified as oncology or non-oncology targets based on parent terms in Disease Ontology. If a protein was targeted by at least 1 oncology drug, it was considered an oncology target.

### Meta-data

Example compounds with exact K_i_ or IC_50_ activity values against human proteins, measured in assays with direct interaction and the highest confidence score=9, were retrieved from ChEMBL v23^65^. Mapping from ENSEMBL identifiers to PDB and polyadenylation data were obtained from GENECODE consortium^67^ version 27. Mapping to enzyme EC numbers, Uniprot and NCBI Gene (Entrez) identifiers were extracted from the HGNC non-alternative loci data set^64^. Target Development Level (TDL) was retrieved from TCRD version 4.6.2^42^. Subcellular localization and protein family information were obtained from UniProt/SwissProt^68^. Probabilities of being loss-of-function intolerant (pLI) were retrieved from Supplementary Data of the ExAC consortium flagship publication^27^. Associations with Mendelian diseases were retrieved from OMIM Morbid Map^32^ (copyright John Hopkins University, AstraZeneca purchased license JHU agreement number A30699 and reference number C03746). We included only binary indicator variables (has/does not have an entry in the Morbid Map). Number of PubMed-indexed articles linked to each gene was retrieved from NCBI Gene^69^ https://www.ncbi.nlm.nih.gov/gene/ on the 02.01.2018. Human to mouse orthologs, K_a_/K_s_ ratios and percentages of sequence identity and similarity were extracted from ENSEMBL Compara^70^ version 91. The lists of essential genes^38^, PTVesc^33^ and rhLOF^39^ genes were obtained from supplementary data of the respective publications. Biological function of the genes was described according to the NCBI Gene/RefSeq summary^71^ unless explicitly indicated otherwise.

### Network analysis

Human protein-protein interaction network was downloaded from STRING v 10.5 (file 9606.protein.links.detailed.v10.5.tsv)^36^. Topological properties were calculated with igraph^72^ version 1.2.1. Weighted k-shell decomposition was computed as described in^73^. Combined evidence scores were used as edge weights for strength, eigenvector centrality and k-shell calculations, i.e., the overall ‘influence’ of a node was proportional to the number of its neighbors combined with confidence in its PPIs. Edge weights were taken as (1 – combined evidence score) for centrality measures based on shortest paths, i.e., shortest paths were the ‘least uncertainty paths’.

### Mapping from GWAS to candidate genes

Genetic associations were obtained from GWAS Catalog^74^ (data set: all associations v1.0, access: 30.08.2017, https://www.ebi.ac.uk/gwas/). Coordinates of genetic variants (SNPs) were mapped from genome assembly GRCh38 to GRCh37 by SNP identifiers (rsids) in 1000 genomes phase 3^75^. Proxy SNPs with r2 >= 0.8 were identified within 50 kilobasepairs in the CEU population using --*hap*-*r2*-positions command with vcftools^76^ version 0.1.13. GWAS variants and their proxy SNPs were mapped to significant single-tissue eQTL from GTEx^30^ version 7.

### Statistical analysis

We applied Fisher exact test for count data because sample sizes were small in some instances (e.g., 3 tissue-specific targets in phase 1 at x = 6) and to be consistent in other analyses. Mann-Whitney U test was used to test differences between groups for continuous variables. Wilcoxon test with explicit handling of tied values in exactRankTests^77^ version 0.8-29 was used to test differences in year of first approval in Fig. 3c. Tests for enrichment or depletion were one-tailed, other tests were two-tailed. Bonferroni correction for multiple testing was applied as appropriate. P-values < 0.05 were considered significant. Statistical analyses were summarized in **Supplementary Information 1**. Figures were generated with ggplot2^78^ version 3.0.0, viridis^79^ version 0.5.1, VennDiagram^80^ version 1.6.20 and UpSetR^81^ version 1.3.3. Analyses were performed in R^82^ version 3.4.1.

### Data availability statement

All data, that were generated in this study, are provided as Supplementary data sets. Annotated Z-score tables for protein-coding genes including the tissue-specific gene and drug target subsets are provided in **Supplementary Data 1**. Network topology properties are provided in **Supplementary Data 2**. Long non-coding RNAs are listed in **Supplementary Data 3**. Columns, that were used as input data for figures, are labelled within each supplementary data set. Summary-level data (counts and percentages) behind figures are included in the **Supplementary Information 1.** Source data for Fig. 4a can be retrieved directly from Ensembl Compara^70^ v 91.

## Author contribution statement

Conceptualization: MR, MH. Formal data analysis: MR. Writing: MR, MH.

## Conflicting interest

MR is a contractor to AstraZeneca. MH is employed by AstraZeneca. AstraZeneca provided support to the authors in form of salaries, but had no role in conceptualization of the study, data collection, analysis, interpretation and writing.

